# Transcriptome remodeling contributes to epidemic disease caused by the human pathogen *Streptococcus pyogenes*

**DOI:** 10.1101/043877

**Authors:** Stephen B. Beres, Priyanka Kachroo, Waleed Nasser, Randall J. Olsen, Luchang Zhu, Anthony R. Flores, Ivan de la Riva, Jesus Paez-Mayorga, Francisco E. Jimenez, Concepcion Cantu, Jaana Vuopio, Jari Jalava, Karl G. Kristinsson, Magnus Gottfredsson, Jukka Corander, Nahuel Fittipaldi, Maria Chiara Di Luca, Dezemona Petrelli, Luca A. Vitali, Annessa Raiford, Leslie Jenkins, James M. Musser

**Affiliations:** Center for Molecular and Translational Human Infectious Diseases Research, Department of Pathology and Genomic Medicine, Houston Methodist Research Institute, and Houston Methodist Hospital, Houston, TX 77030; Departments of Pathology and Laboratory Medicine, and Microbiology and Immunology, Weill Cornell Medical College, New York, NY 10021; Section of Infectious Diseases, Department of Pediatrics, Texas Children’s Hospital and Baylor College of Medicine, Houston, TX 77030; Department of Medical Microbiology and Immunology, Medical Faculty, University of Turku, Turku, Finland; Department of Infectious Diseases, National Institute for Health and Welfare, Turku, Finland; Departments of Clinical Microbiology and Infectious Diseases, Landspitali University Hospital, Reykjavik, Iceland; Faculty of Medicine, School of Health Sciences, University of Iceland, Reykjavik, Iceland; Department of Mathematics and Statistics, University of Helsinki, Helsinki, Finland; Public Health Ontario, and Department of Laboratory Medicine and Pathobiology, Faculty of Medicine, University of Toronto, Toronto, Ontario, Canada; School of Pharmacy, University of Camerino, Camerino, Italy; School of Biosciences and Veterinary Medicine, University of Camerino, Camerino, Italy

## Abstract

For over a century, a fundamental objective in infection biology research has been to understand the molecular processes contributing to the origin and perpetuation of epidemics. Divergent hypotheses have emerged concerning the extent to which environmental events or pathogen evolution dominates in these processes. Remarkably few studies bear on this important issue. Based on population pathogenomic analysis of 1200 *Streptococcus pyogenes* type *emm*89 infection isolates, we report that a series of horizontal gene transfer events produced a new pathogenic genotype with increased ability to cause infection, leading to an epidemic wave of disease on at least two continents. In the aggregate, these and other genetic changes substantially remodeled the transcriptomes of the evolved progeny, causing extensive differential expression of virulence genes and altered pathogen - host interaction, including enhanced immune evasion. Our findings delineate the precise molecular genetic changes that occurred and enhance our understanding of the evolutionary processes that contribute to the emergence and persistence of epidemically successful pathogen clones. The data have significant implications for understanding bacterial epidemics and translational research efforts to blunt their detrimental effects.

**Importance:** The confluence of studies of molecular events underlying pathogen strain emergence, evolutionary genetic processes mediating altered virulence, and epidemics is in its infancy. Although understanding these events is necessary to develop new or improved strategies to protect health, surprisingly few studies have addressed this issue, in particular at the comprehensive population genomic level. Herein we establish that substantial remodeling of the transcriptome of the human-specific pathogen *Streptococcus pyogenes* by horizontal gene flow and other evolutionary genetic changes is a central factor in precipitating and perpetuating epidemic disease. The data unambiguously show that the key outcome of these molecular events is evolution of a new, more virulent pathogenic genotype. Our findings provide new understanding of epidemic disease.

## Introduction

Genetic diversity begets phenotype variation and with it the possibility of a different life. Considerable effort has been expended in the last 40 years to understand the genetic diversity and population structure of many bacterial pathogens, especially those that detrimentally affect human and livestock health and cause epidemics (1-27). These studies have led to the general concept that some bacterial species are clonal, with relatively little evidence that horizontal gene transfer (HGT) and recombination shape species diversity, whereas other bacterial pathogens are highly recombinogenic, with species diversity mediated by extensive HGT events (1-27). Genetic studies have been greatly facilitated in recent years by relatively inexpensive large-scale comparative DNA sequencing, which now makes it possible to precisely delineate the nature and extent of genomic variation present in large populations (hundreds to many thousands) of individual pathogenic bacterial species (4, 5, 10-18, 23-26). For example, analyses of important pathogens such as *Staphylococcus aureus, Streptococcus pyogenes, Streptococcus pneumoniae, Escherichia coli, Salmonella enterica* serovars, and *Legionella pneumophila* have been conducted, resulting in much new information about genetic variation in these and other species (4, 5, 10-18, 23-26, 28-35).

In parallel with studies of bacterial population genetic structure, there has been interest in identifying the precise genomic changes that contribute to the emergence, numerical success, and epidemic behavior of members of some bacterial species. A major effort has been devoted to analysis of comprehensive, population-based samples of the strict human pathogen *S. pyogenes* (commonly, group A streptococcus or GAS) as a model pathogen (28-35). *S. pyogenes* is endemic in humans worldwide and periodically causes epidemics of superficial (e.g. pharyngitis and impetigo) and invasive (e.g. necrotizing fasciitis, pneumonia, myositis) infections. Globally, the organism causes an estimated 711 million human infections and over 500,000 deaths annually (36). The species is genetically diverse, with more than 240 *emm* gene types (www.cdc.gov/abcs/index.html) and approximately 650 multilocus sequence types (MLSTs) (spyogenes.mlst.net) described.

In the early 1980s a dramatic increase in the frequency and severity of infections caused by *S. pyogenes* led to the recognition of a global pandemic caused by *emm*1 strains (37-44). This pandemic afforded the opportunity to compare pre-epidemic and epidemic strains for potential bacterial factors contributing to this global health problem. To gain insight into the emergence, dissemination, and diversification of *emm*1 strains causing this pandemic, we sequenced the genome of 3,615 *emm*1 infection isolates (32). Phylogenetic analyses revealed that the emerged pandemic *emm*1 strains are a genetically closely related clonal population that evolved from its most recent pre-epidemic progenitor in the early 1980s. The key genetic event underpinning the pandemic was acquisition by HGT and recombinational replacement of a 36-kb segment of the *S. pyogenes* core chromosome that mediated enhanced production of toxins NAD+glycohydrolase (SPN, *S. pyogenes* NADase) and streptolysin O (SLO) (32). A subsequent study (35) showed that the striking upregulation of SPN and SLO production by members of the pandemic clone and altered virulence phenotype occurred as a consequence of only three single nucleotide polymorphisms (SNPs). Two are located in the ‐35 to ‐10 spacer region of the promoter sequence upstream of the *nga-ifs-slo* transcriptional unit and resulted in increased gene expression. The third, a nonsynonymous SNP in the *nga* gene, increases the activity of SPN, a secreted cytotoxin virulence factor (45). Additional evidence supporting upregulation of SPN and SLO as a contributing cause of *S. pyogenes* epidemic disease was found by sequence analysis of 1,125 *emm*89 genomes (35) obtained in comprehensive population-based surveillance studies conducted in the United States, Finland, and Iceland between 1995 and 2013. Among these *emm*89 strains we identified three distinct phylogenetic clades (designated clade 1, clade 2, and clade 3). The current worldwide recent increase in *emm*89 invasive infections corresponded temporally with the emergence and expansion of clade 3 strains upregulated in SPN and SLO production (35, 46).

Thus, progress is being made in understanding genomic alterations that are linked with increases in disease frequency and severity in some human pathogens. However, despite these advances, very little analogous work has been conducted to investigate global changes in gene expression that may contribute to the origin and perpetuation of bacterial epidemics. Similarly, there is a general lack of studies linking genome variation, transcriptional changes, and altered virulence in epidemic forms. The primary goal of this investigation was to study how genome variation linked with changes in transcriptome and altered virulence might contribute to the origin and perpetuation of bacterial epidemics, using the ongoing *S. pyogenes *emm**89 epidemic as a convenient model system. We used comparative pathogenomics to dissect the precise molecular genetic events that have mediated the evolutionary origin and diversification of the epidemic *emm*89 strains. Unexpectedly, we found that a high frequency of HGT events has shaped the *emm*89 population genetic structure to a far greater extent than vertically inherited SNPs and short insertions and deletions (indels). Global transcriptome (RNAseq) analysis was conducted on genetically representative pre-epidemic and epidemic *emm*89 strains to determine the extent to which the genomic changes causing altered gene expression may have contributed to the epidemic. We found that HGT is extensive in the *emm*89 population and has contributed disproportionately to the diversification of virulence factors and their expression. Nonsynonymous SNPs in major regulatory genes and other modest genetic changes have also led to transcriptome remodeling intimately linked with the origination and perpetuation of the epidemic. The results have significant implications for understanding epidemic bacterial disease and translational research efforts designed to control or limit the detrimental effect of infectious agents. The overall strategy used herein is of general utility and pertinence to the investigation of other pathogens.

## Results and Discussion

### Population Genetic Structure and Contribution of Horizontal Gene Transfer (HGT)

We studied 1,200 *emm*89 *S. pyogenes* strains, virtually all (*n*=1,198) cultured from patients with invasive infections that occurred between 1995 and 2014 (Fig. 1, Table S1). The great majority of strains (*n*=1,180) were collected as part of comprehensive population-based studies conducted in the United States, Finland, and Iceland. The genomes of all 1,200 strains were sequenced to a mean 60-fold depth of coverage (range 13-to-440) using an Illumina paired-end strategy, and polymorphisms were identified. Inference of genetic relationships and Bayesian clustering showed that these *emm*89 strains have a major population of 1,193 strains comprising 3 primary genetic clades (Fig. 2). Seven substantially divergent *emm*89 strains are genetic outliers (Fig. 3). The genomes of 3 strains representing the genetic backgrounds of organisms assigned to the three primary clades (strains MGAS11027, MGAS23530, and MGAS27061 respectively) were closed and annotated (Fig. S1). The epidemiological information available for the 1,200 strains revealed that clade 3 strains emerged and expanded rapidly in the United States, Finland and Iceland, displacing their predecessor clade 1 and 2 strains in the populations studied (Fig. 1). These findings are consistent with the preliminary data that we recently reported (35).

**Fig. 1.**
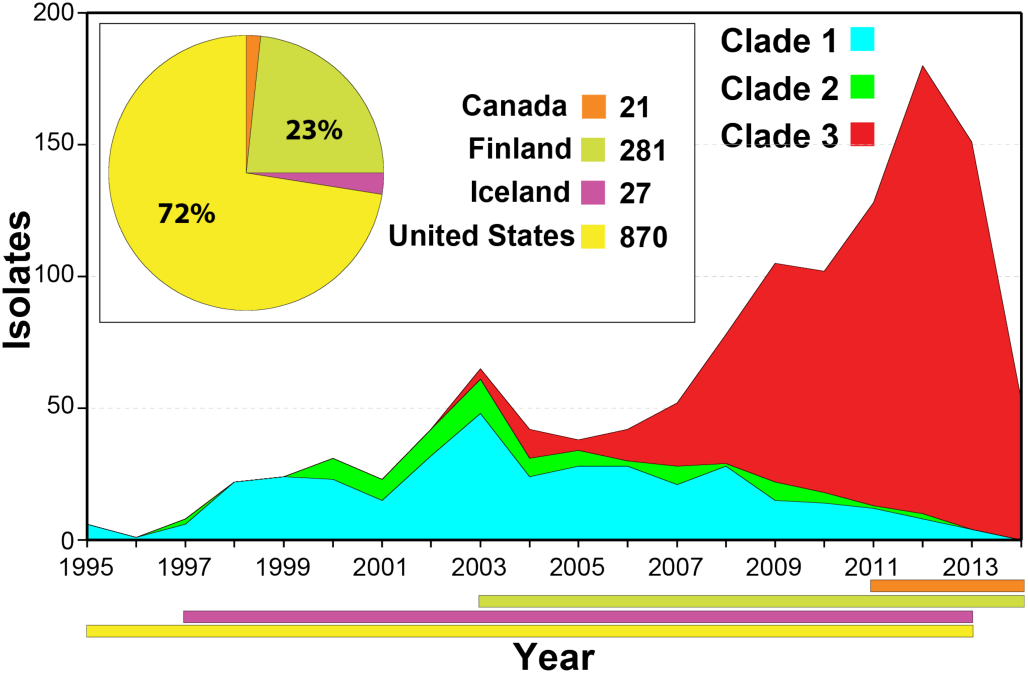
Temporal and geographic distribution of the *emm*89 strain cohort. Shown is the temporal distribution of the *emm*89 strains by clade. The inset contains the geographic distribution of the isolates by country. The colored horizontal bars at the bottom of the figure show the temporal distribution of the strains by country. A single isolate from Italy is not illustrated. The reduced cases in 2014 are due to U.S. isolates not being available for study, not a decline in the frequency of infections. Clade 3 strains emerged in 2003 and expanded greatly, displacing predecessor clade 1 and 2 strains in all 3 populationbased strain samples studied (United States, Finland, and Iceland).

**Fig. 2.**
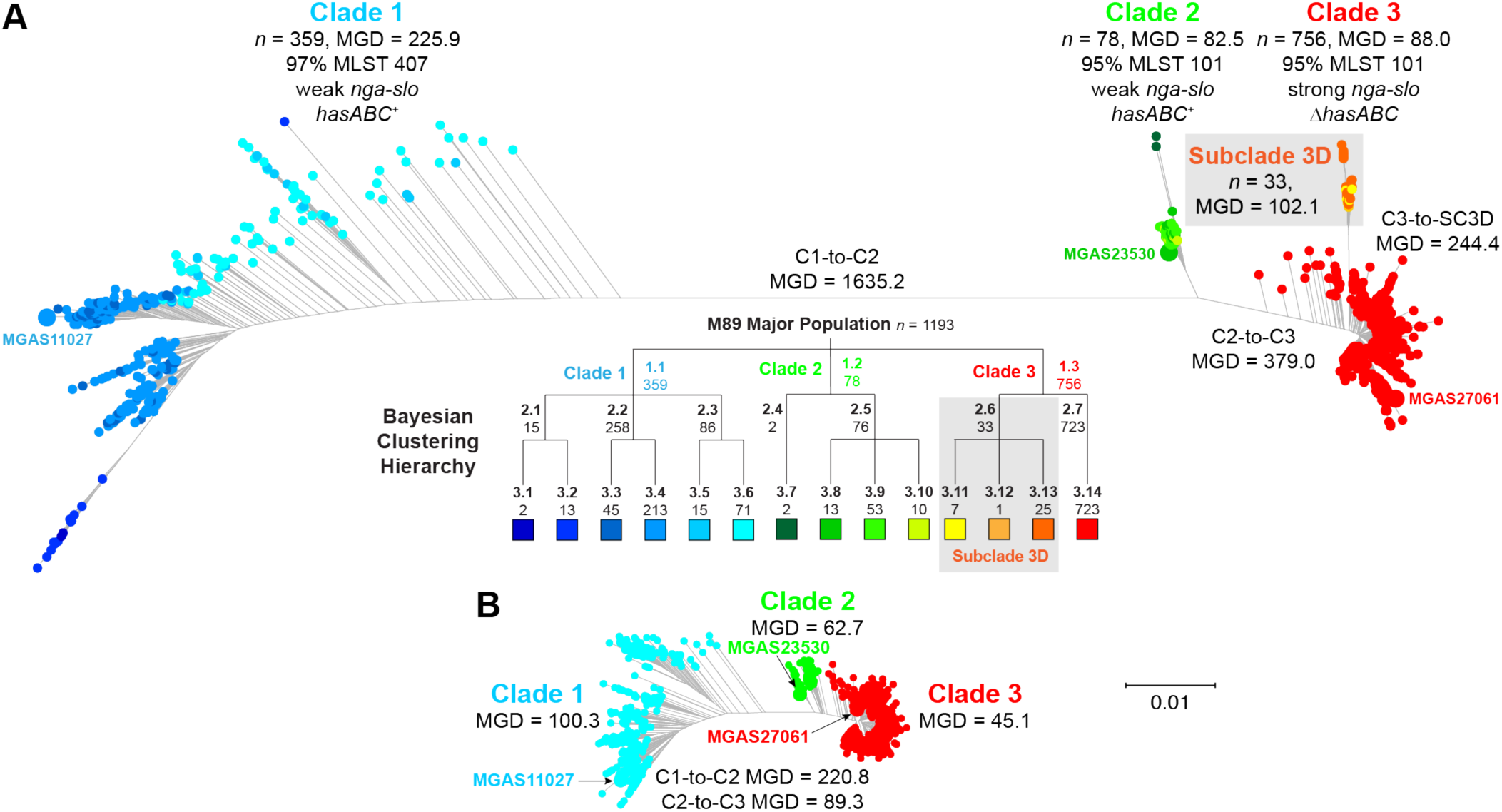
Genetic relationships among the major population of *emm*89 strains. Genetic relationships were inferred by the neighbor-joining method based on concatenated core chromosomal SNPs using SplitsTree. Indicated for the inferred phylogenies are the mean genetic distances (MGDs) both inter‐ and intra-clade measured as differences in core chromosomal SNPs. (*A*) Inferred genetic relationships based on 11,846 SNPs identified among the major population 1,193 strains. Isolates are colored by cluster as determined using BAPS as indicated in the hierarchy below the figure. Three major clades (C1, C2, and C3) are defined at the first level of clustering. Subclade 3D (SC3D), a recently emerged and expanding population of strains in Finland, is defined at the second level of clustering. The mean genetic distance among strains within clades is less than the MGD to strains of the nearest neighboring clade(s). Bootstrap analysis with 100 iterations gives 100% confidence for all of the clade-to-clade branches (i.e. C1-C2, C2-C3, and C3-SC3D). (*B*) Genetic relationships inferred based on 8,989 SNPs identified among the major population of 1,193 strains, filtered to exclude horizontally acquired sites as inferred using GUBBINs. Exclusion of sites attributed to horizontal gene transfer events collapses the MGD strain-to-strain both within and between the clades. The MGD within the clades remains less than the MGD to the nearest neighboring clade(s). Both trees are illustrated at the same scale.

**Fig. 3.**
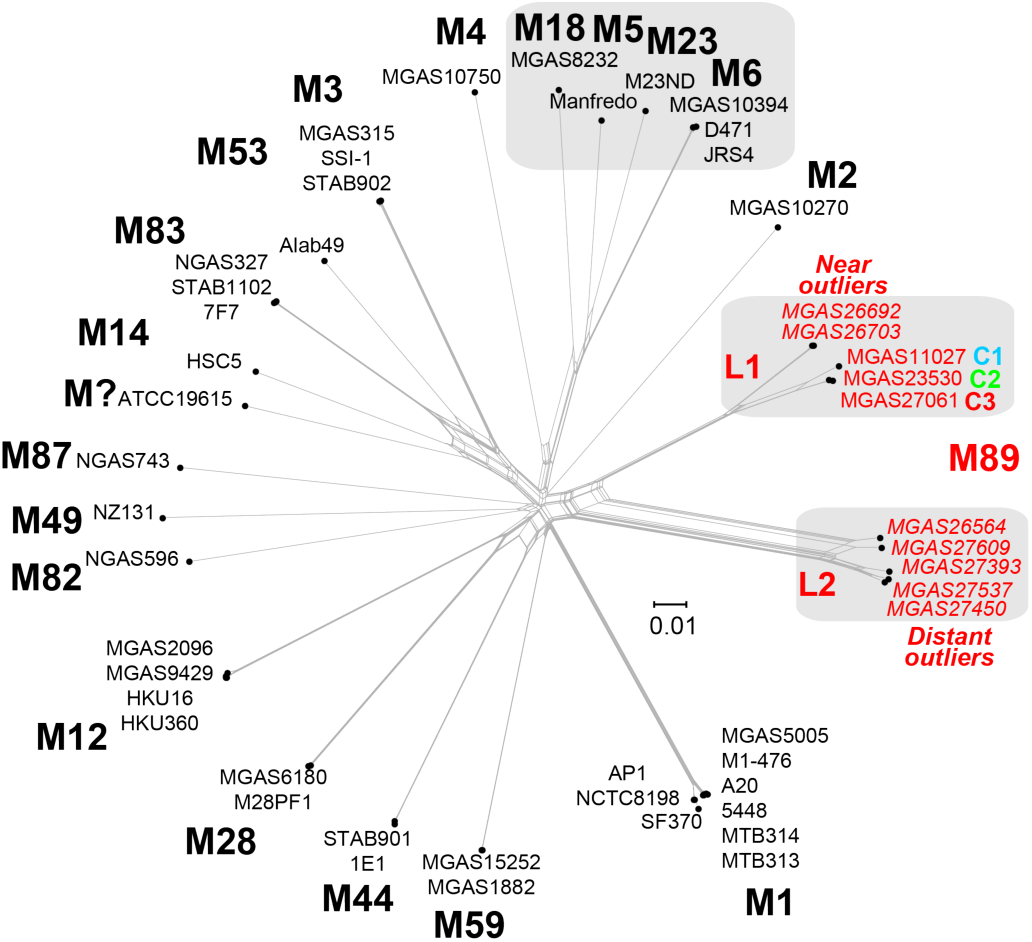
Genetic relationships between strains of various Emm/M protein serotypes. Genetic relationships were inferred among 49 GAS strains of 20 M-types based on 75,184 concatenated core chromosomal SNPs by the neighbor network method. The analysis is based on 42 closed genomes and 7 whole-genome-sequenced *emm*89 genetic outlier strains shown in italics. The MGD inter-serotype is 16,340 SNPs. *emm*89 strains are the only *emm* type with two distinct lineages (L1 and L2) in the interserotype network. The MGD of 14,247 SNPs between the *emm*89 L1 and L2 genomes is greater than the MGD of 11,548 SNPs among the serotype M5, M6, M18, and M23 genomes. Of note, the *emm*89 L1 to L2 MGD is greater than the *emm*89 L1 to M53 genome MGD of 14,194 SNPs.

The *emm*89 population genomic data revealed an unprecedented level of genetic diversity for strains of a single *S. pyogenes emm* type. Comparison of the *emm*89 genome sequences with data available for 37 *S. pyogenes* genomes of 18 other *emm*-types (Table S2) showed the *emm*89 strains are the only *emm*-type to have two deeply rooted branches in the phylogenetic network (Fig. 3).

We identified extensive genomic diversity between and within the three primary *emm*89 clades. The mean genetic distance (MGD) among the 1,193 strains of the 3 clades was 610 SNPs in the core genome (Fig. 2A). In striking contrast, among 3,615 *emm*1 strains collected in 8 countries on two continents over 45 years (*i.e*., a collection 3 times larger, from a broader geographic region, and a 2.5 times longer period than the *emm*89 sample) the MGD was only 106 core SNPs (32).

There was a nonrandom distribution of SNPs throughout the *emm*89 genomes. Multiple regions had elevated SNP density, indicating HGT and genomes with a mosaic evolutionary history (Fig. 4, Fig. S1). GUBBINs statistical analysis of SNP distribution (47) identified 2,316 regions of putative HGT with a mean size of 3,695 bp (range, 4 bp to 71,774 bp) at 526 loci around the genome. Because HGT can distort inferences of genetic relationships and evolutionary history, the phylogeny of the strains was reassessed using sequences filtered to exclude regions of recombination (Fig. 2B). This analysis greatly reduced the MGD (i.e. average pairwise core SNPs) among the 1,193 strains by 78%, from 610 to 134, a level similar to that found in 3,615 *emm*1 strains. The MGD from clade 1 to clade 2, and from clade 2 to clade 3 was reduced by 87% and 76%, respectively. However, importantly, 3 distinct clades remained among the 1,193 strains.

**Fig. 4.**
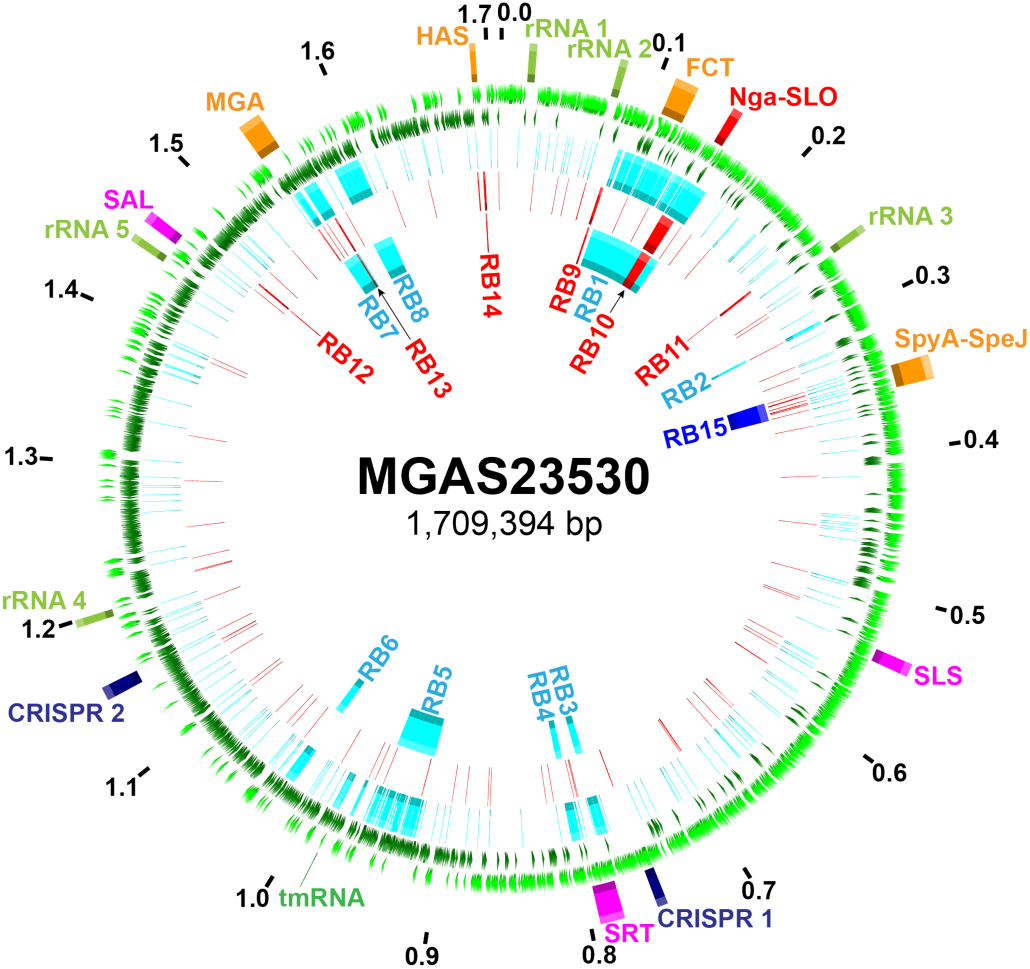
Distribution of SNPs and regions of horizontal gene transfer. Illustrated in the genome atlas of clade 2 strain MGAS23530 from the 1st (outer-most) ring to the 7th (inner-most) ring are as follows. 1) Genome size in megabase pairs (black). 2) Landmarks: rRNA, 23S-16S-5S ribosomal RNA; FCT, fibronectin/collagen/T-antigen; SLS, streptolysin S; SRT, streptin; SAL, salvaricin; MGA, mga operon; HAS, *hasABC* capsule synthesis operon. 3 & 4) Coding sequences on the forward (light-green) and reverse (dark-green) strands. 5) Clade 1 strain MGAS11027 SNPs (*n*=1,915, light-blue) relative to clade 2 strain MGAS23530. 6) Clade 3 strain MGAS27061 SNPs (*n*=415, red) relative to clade 2 strain MGAS23530. 7) Predicted regions of horizontal gene transfer separating clade 1 and 2 strains (lightblue), clade 2 and 3 strains (red), and clade 3 and subclade 3D strains (dark-blue) as listed in Table 1. SNPs are nonrandomly distributed. Regions of elevated SNP density correspond to predicted horizontal gene transfer/recombination blocks.

Outgroup rooting with the genome of *emm*1 reference strain SF370 showed that the evolutionary pathway leading to the current *emm*89 epidemic lineage had clades branching in the sequence of clade 1, followed by clade 2, and then clade 3 (Fig. S2). Clade 1 and clade 2 strains differed by 8 regions of HGT involving 171.1 kb (10% of the genome), and clade 2 and clade 3 strains differed by 6 regions of HGT (15.3 kb, 0.9% of the genome) (Fig. 4, Table 1). Seven of the 8 HGT regions differentiating clade 1 and clade 2 are most similar in sequence to regions in *emm*2 reference genome MGAS10270 (Fig. S3). Of special note, 33 isolates in clade 3 differed from the 725 other clade 3 strains by one additional HGT (Fig. 2A). These strains, designated subclade 3D (SC-3D) (Fig. 2A), first occur in the Finland sample in 2009, and have disproportionately increased in recent years as a cause of bloodstream infections in that country (Table S1, Fig. S4).

**Table 1.** HGT recombination blocks separating GAS *emm89*/M89 clades.

| Block | Clades | Start<sup>a</sup> | Stop<sup>a</sup> | Length | SNPs | Genes | M-Like | %ID |
| --- | --- | --- | --- | --- | --- | --- | --- | --- |
| RB1<sup>b</sup> | C1-C2 | 92,389 | 164,162 | 71,774 | 411 | 72 | M2 | 88.74<sup>b</sup> |
| RB2 | C1-C2 | 295,481 | 297,574 | 2,094 | 9 | 2 | M2 | 100.00 |
| RB3<sup>c</sup> | C1-C2 | 773,487 | 780,634 | 7,148 | 55 | 8 | M2 | 99.55 |
| RB4<sup>c</sup> | C1-C2 | 794,417 | 800,659 | 6,243 | 28 | 8 | M2 | 99.55 |
| RB5 | C1-C2 | 921,261 | 960,297 | 39,037 | 100 | 41 | M2 | 99.68 |
| RB6 | C1-C2 | 1,022,619 | 1,030,407 | 7,789 | 20 | 6 | M2 | 99.97 |
| RB7 | C1-C2 | 1,543,651 | 1,561,165 | 17,515 | 103 | 13 | M2 | 97.17 |
| RB8 | C1-C2 | 1,577,916 | 1,597,447 | 19,532 | 138 | 21 | M28 | 99.58 |
| RB9 | C2-C3 | 86,603 | 88,366 | 1,764 | 12 | 2 | M5/M23 | 99.38 |
| RB10 | C2-C3 | 145,163 | 155,569 | 10,407 | 59 | 11 | M1/M12 | 98.52 |
| RB11 | C2-C3 | 244,407 | 244,758 | 352 | 5 | 1 | M5 | 100.00 |
| RB12 | C2-C3 | 1,472,262 | 1,473,025 | 764 | 9 | 1 | M12 | 100.00 |
| RB13 | C2-C3 | 1,558,898 | 1,559,698 | 801 | 7 | 2 | M49 | 99.75 |
| RB14 | C2-C3 | 1,693,613 | 1,694,805 | 1,193 | 6 | 2 | M5/M6 | 100.00 |
| RB15 | C3-SC3D | 341,762 | 359,579 | 17,818 | 106 | 21 | M1 | 99.67 |

<sup>a</sup> Start and stop positions provided are relative to the MGAS23530 genome.
<sup>b</sup> The first 18.6 kb and last 39.6 kb are M2-like (>99% ID), however the central 13.5 kb FCT pilus encoding region is unlike that of any other sequenced GAS *emm*-type.
<sup>c</sup> RB3 and RB4, likely represent a single HGT event that encompasses the intervening streptin lantibiotic synthesis genes, thus resulting in a larger single recombination of 26,697 bp.

HGT events are responsible for the bulk of the sequence difference between the clades. The transferred sequences encompass multiple genes encoding many known secreted and cell surface-associated virulence factors, including the pilus/T-antigen adhesin, fibronectin-binding protein FbaB, the toxin pair NGA and SLO, internalin InlA, C5a peptidase ScpA, antiphagocytic M-like proteins Enn and Mrp, virulence regulators Mga and Ihk-Irr, immunogenic secreted protein Isp1, and the capsule synthesis enzymes HasABC (47). These HGT events have had important consequences. For example, clade 1 strains differ from clade 2 and 3 strains in pilus/T-antigen, and the clade 3 strains cannot produce capsule due to loss of the *hasABC* genes. Of note, different pilus types have been shown to vary in cell adherence and tissue tropism, and differences in the level of production of capsule and SPN and SLO cytotoxins can alter virulence (35, 47, 48).

Consistent with SPN and SLO playing a key role in *S. pyogenes* strain emergence and enhanced fitness, each of the three clades has a distinct *nga-ifs-slo* region resulting from two independent HGT events. In addition, SC-3D strains differ from the other clade 3 strains due to HGT of a region encoding the SpyA and SpeJ virulence factors (47, 49-51). Inasmuch as these multiple HGT events involve regions encoding virulence factors, it is reasonable to hypothesize that many of these HGT events alter host-pathogen interaction.

### Variation in Gene Content and Phage Genotypes

HGT in bacteria can be mediated by mobile genetic elements, phages and integrative-conjugative-elements (ICEs). *S. pyogenes* phages commonly encode one or more secreted virulence factors such as streptococcal pyrogenic exotoxin superantigens and streptococcal phage DNases (52, 53). *S. pyogenes* ICEs usually encode one or more factors mediating resistance to antibiotics such as tetracycline and macrolides (52). Horizontal acquisition of antibiotic resistance and novel virulence factor genes, mediated by ICEs and phages, has been associated with localized outbreaks and large epidemics of *S. pyogenes* infections (29). Mobile genetic element (MGE) content was investigated in 1,193 *emm*89 isolates relative to the combined gene content (> 53,000 genes) of 30 GAS genomes of 18 *emm*-types (Tables S1 and S2). This analysis identified 64 different profiles of MGE content (Fig. 5). ICEs were infrequent in the strain sample. The 3 most prevalent MGE profiles accounted for 72% of the strains (phage genotypes (PGs) PG01, PG02 and PG03), corresponding with the phage content of the reference genomes for each of the three clades (Fig. 5, Fig. S1). With the exception of PG02 (defined as lack of prophages), most phage genotypes were confined to a single clade. The most prevalent PG in clade 1 was PG03 (43%; phages 11027.1 encoding SpeC and Spd1, and 11027.2 encoding Sdn). Also prevalent were PG05 (13%) and PG06 (11%) strains, potentially derived from PG03 strains by phage loss. Most clade 2 strains are PG02 (72%), having no phages. The abundance of PG02 strains representing 20% of the entire *emm*89 cohort is unusual in that prior to our investigation nearly all *S. pyogenes* genomes have been found to be polylysogenic (53). Most clade 3 strains are PG01 (62%), having phage 27061.1 encoding SpeC and Spd1, followed next in prevalence by PG02 (22%). Of note, although phages 11027.1 and 27061.1 are integrated at the same genomic locus and encode the same two secreted virulence factors, they are different phages (Fig. S5). PG01 (presence of 27061.1) first occurred in our strain samples in 2003, a time that corresponds with the emergence of the epidemic clade 3 strains. However, the acquisition of 27061.1 into the *emm*89 population does not result in the epidemic clade 3 strains acquiring new phage-encoded virulence genes that were not already prevalent in the pre-epidemic clade 1 strains. This finding suggests that acquisition of phage-encoded virulence genes was not a key driver for the emergence of epidemic clade 3 organisms as has been speculated (46).

**Fig. 5.**
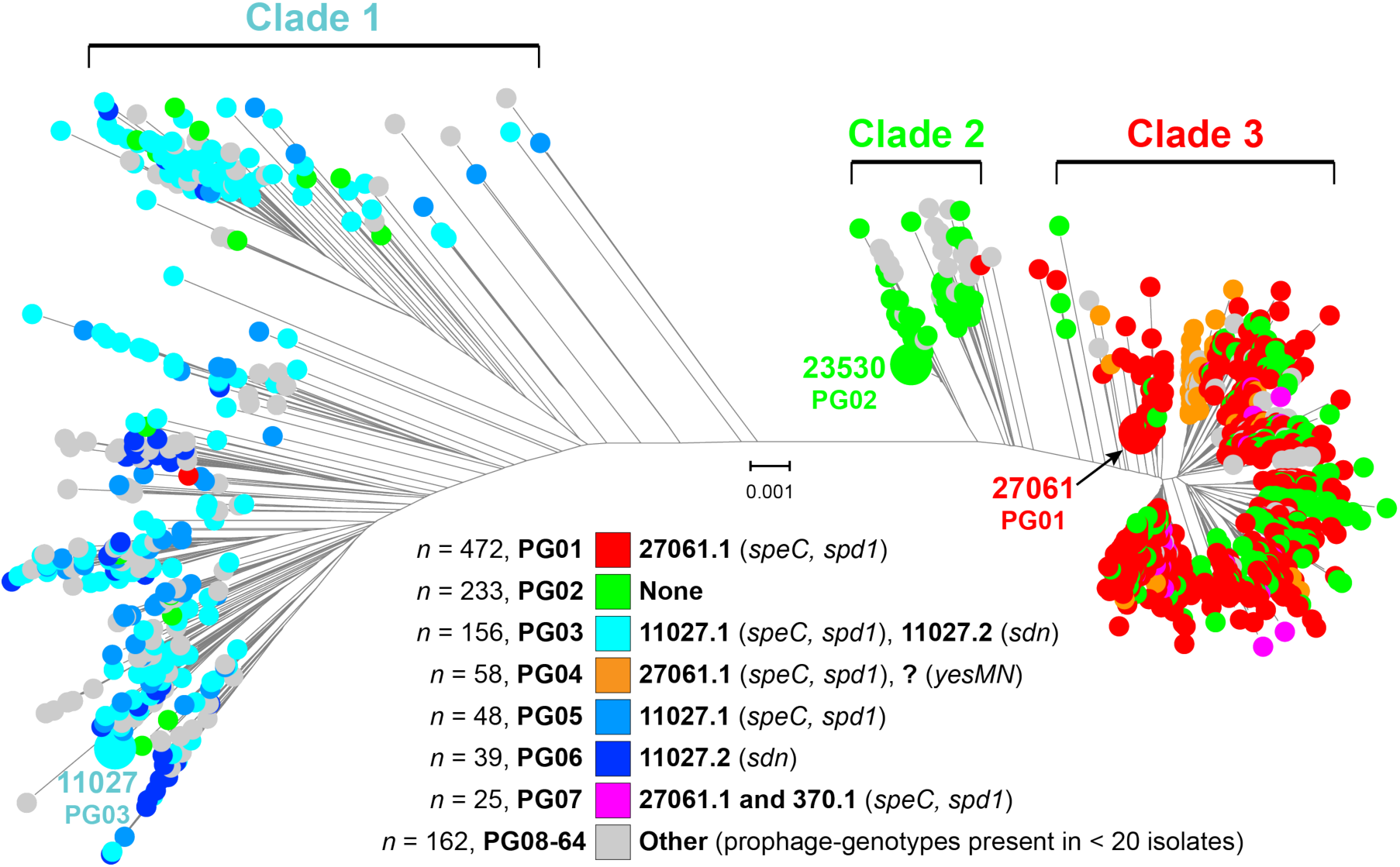
Prophage content of the *emm*89 strains. Shown is the phylogeny inferred by neighbor-joining for the 1,193 clade 1, 2, and 3 isolates based on 8,989 core SNPs filtered to exclude SNPs acquired by horizontal gene transfer events. Isolates are colored by phage genotype (PG) as indicated in the index. PGs were assigned in order of prevalence of occurrence in the strain sample. With the exception of PG02 (absence of phage), most of the PGs are exclusive to a single clade. PG01 is first present in the strain sample in 2003 in two isolates, one each of clade 2 and 3. 2003 is also when epidemic clade 3 strains are first present in the strain sample.

### HGT and Extensively Remodeled Global Transcriptomes

One school of thought postulates that HGT events are similar to point mutations in that most of them are neutral or nearly so, and have little effect on pathogen traits. The unexpected magnitude of HGT events in the study population (based on previous findings from analysis of other *S. pyogenes emm* types) provided a unique opportunity to test the hypothesis that these HGT events have enhanced the virulence of the epidemic *emm*89 strains by remodeling of the global transcriptome. As a consequence of its greater technical difficulty and expense, global transcriptional variation has been far less studied than genomic variation in bacterial pathogens. Moreover, since the samples studied herein are population-based, comprehensive, and include temporal-spatial information, we had the additional opportunity to assess the potential effect of transcriptome remodeling on strain emergence and dissemination. We used RNAseq to compare transcript variation at two growth points among genetically representative strains of clades 1, 2, and 3 (Fig. 6). The strains lack polymorphisms in major regulatory genes such as *covRS, mga,* and *ropB* known to influence *S. pyogenes* gene expression and virulence (47, 54-58). The number of genes differentially expressed in stationary-phase growth exceeded the number in exponential-phase growth by approximately 3-fold in all of the clade-to-clade comparisons (Fig. 7A). A general finding was that the greater the genetic distance between strains, the greater the number of genes significantly altered in transcription. The largest number of differentially expressed genes was recorded between strains MGAS11027 (clade 1) and MGAS23530 (clade 2), consistent with strains in these clades being separated by the greatest MGD (Fig. 2). Genes altered in transcript level by 1.5-fold or greater accounted for 14% and 36% of the genome at exponential and stationary growth phases, respectively, in comparing MGAS11027 (clade 1) and MGAS23530 (clade 2) (Table S3 section 1). Although genes (*n*=182) located within the eight distinct regions of HGT differentiating clade 1 and clade 2 comprise only 11% of the genome, at exponential growth they accounted for 24% of the differentially expressed genes, a highly nonrandom occurrence (P<0.0001). Importantly, genes encoding many key virulence factors had significantly different transcript levels, including the FCT region pilin genes, *nga-ifs-slo, speG, ideS, ska, sclA, fba, enn, emm, mrp, and mga* (47). Collectively, these findings demonstrate that the genome segments that have been horizontally acquired and retained on the evolutionary pathway leading from clade 1-to-clade 2 strains have contributed disproportionately to remodeling the global transcriptome, including many virulence genes, and argue that they are likely not selectively neutral.

**Fig. 6.**
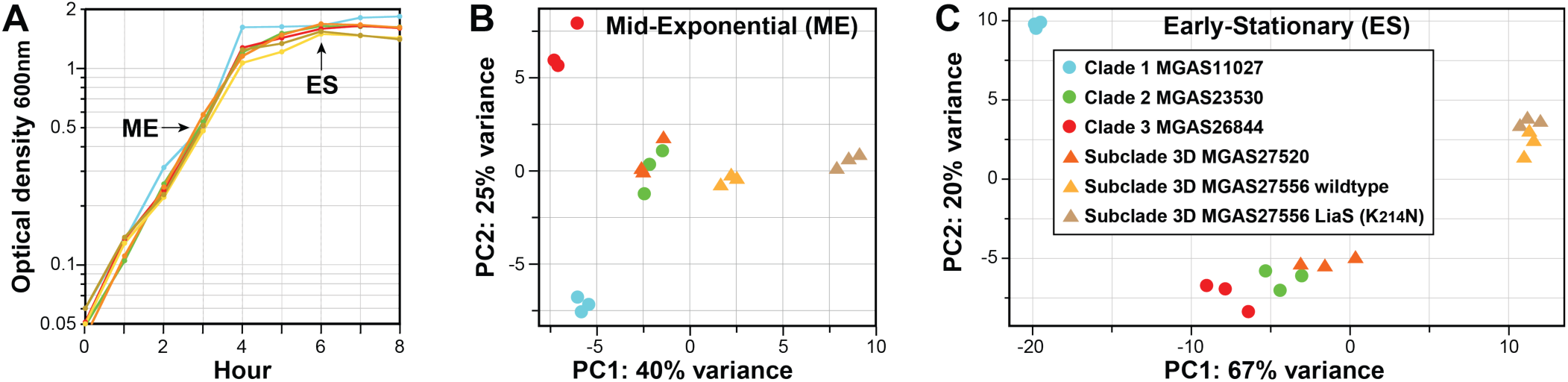
Transcriptome analysis of genetically representative pre-epidemic and epidemic *emm*89 strains. RNAseq analysis was done in triplicate for six genetically representative strains. The strain index provided in panel C applies to all of the panels. (*A*) Growth curves. Graphed is the average of growth curves done in triplicate. The growth curves were closely similar for all strains. Cells were harvested for RNA isolation at mid-exponential (ME = OD_600_ 0.5) and early stationary growth (ES = 2 hr postexponential phase). (*B & C*) Principal component analyses. Illustrated are transcriptional variances among the strains expressed as the primary and secondary principal components, the two largest unrelated variances in the data. Strain replicates cluster together, illustrating good reproducibility.

**Fig. 7.**
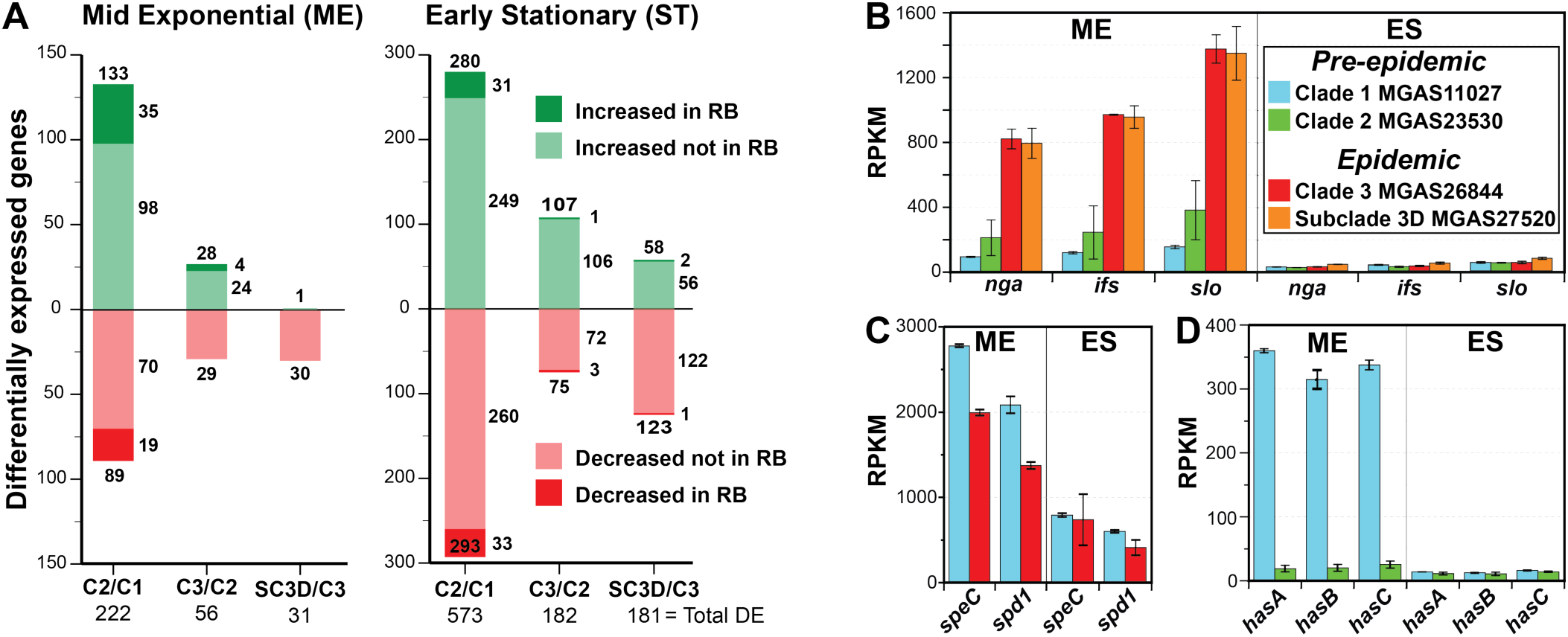
RNAseq expression analysis. (*A*) Genes significantly differentially expressed (DE) at 1.5-fold change or greater. Indicated for each comparison is the total number of differentially expressed genes both within and outside of the recombination blocks (RBs). The representative strains of each clade analyzed are: C2/C1 = MGAS23530/MGAS11027, C3/C2 = MGAS26844/MGAS23530, and SC-3D/C3 = MGAS27520/MGAS26844. (*B*) Transcript levels for the *nga-ifs-slo* operon. The transcript levels of *nga*, *ifs*, and *slo* are significantly greater in the epidemic strains than the pre-epidemic strains by 4‐ to 8fold (P<0.05) at mid-exponential growth. The index in panel B applies to panels B, C and D. (*C*) Transcript levels for the phage-encoded virulence factors *speC* and *spd1.* The transcript levels of *speC* and *spd*1 are significantly greater in the pre-epidemic strain at mid-exponential growth (P<0.01). (*D*) Transcript levels for the *hasABC* operon. Transcription of *hasABC* is very weak for the clade 2 strain at both growth phases, and is significantly less than for the clade 1 strain at mid-exponential growth (P<0.002). RB, recombination block; RPKM, reads per kilobase per million reads mapped.

The genomic changes accruing in the molecular evolution of clade 2-to-clade 3 are of considerable interest because they are associated with the emergence, dissemination and recent rapid increase in frequency of *emm*89 invasive infections recorded in many countries (46, 59-63). We found that 1% of the genome was reshaped in the clade 2-to-clade 3 transition, a much more modest number than for the clade1-to-2 transition. However, differentially expressed genes accounted for 4% and 11% of the genome at exponential and stationary growth phases, respectively, in comparing the two representative strains MGAS23530 (clade 2) and MGAS26844 (clade 3) (Table S3 section 2, Fig. 7A). Genes located within regions of HGT were significantly overrepresented among the differentially expressed genes in exponential growth (P<0.0001). Included among the 28 genes with significantly increased expression in exponential growth were the critical virulence genes *nga-ifs-slo* (Fig. 7B). Importantly, significantly increased transcription of *nga-ifs-slo* was associated with the emergence and epidemic increase in *S. pyogenes emm*1 invasive infections (32, 34, 35).

Additional genetic changes that differentiate epidemic clade 3 strains from the most recent predecessor clade 2 strains are acquisition of phage 27061.1 encoding *speC* and *spd*1 and loss of the *hasABC* capsule synthesis genes. To explore the potential role these genetic changes have played in contributing to the emergence of the epidemic clade 3 strains, we inspected transcript data for the *speC* and *spd1,* and *hasABC* virulence factor genes between the pre-epidemic (clade 1 and 2) and epidemic (clade 3) *emm*89 representative strains. Transcript levels of *speC* and *spd*1 were significantly greater for the pre-epidemic clade 1 strain MGAS11027 than epidemic clade 3 strain MGAS26844 at both phases of growth assessed (Fig. 7C). The finding of significantly lower *speC* and *spd*1 transcripts in the genetically representative epidemic clade 3 strain further argues that presence of these virulence factors in the clade 3 lineage unlikely confers a fitness advantage relative to clade 1 strains, and therefore is an unlikely mechanism for the emergence of the epidemic clone and displacement of the predecessor clade 1 and clade 2 strains (46). Similarly, although the epidemic clade 3 strains are incapable of producing the antiphagocytic hyaluronic acid (HA) capsule due to HGT-mediated loss of the *hasABC* genes, the transcript data indicate this gene loss was not likely responsible for a significant decrease in capsule production between the clade 2 and 3 strains. We found that transcription of *hasABC* was very weak in clade 2 strain MGAS23530 under both growth conditions assessed (Fig. 7D), arguing that capsule production was already negligible before the HGT mediated loss of the *hasABC* genes by the clade 3 lineage. Capsule production was strong only for the clade 1 strain MGAS11027 at exponential growth.

We next investigated the molecular basis for the difference in capsule production using all strains of clades 1 and 2. Sequence variation in the *hasABC* promoter has been reported to alter transcription and capsule production (64). Inspection of the genome sequence data, coupled with Sanger sequencing of the *hasABC* promoter for all clade 1 and 2 strains identified two major variants (Fig. S6A). These promoter variants corresponded with strong clade 1 strain MGAS11027 and weak clade 2 strain MGAS23530 *hasABC* transcription. Whereas both promoter variants are equally represented among clade 1 strains, the vast majority (88.5%) of clade 2 strains have the weak transcription variant (Fig. S6B and S6C). Thus, the evolution of clade 3 from a clade 2 progenitor strain likely involved transition from very little capsule to no capsule production. This again argues that loss of the *hasABC* genes by the clade 3 lineage is unlikely to confer a fitness advantage relative to the clade 2 strains and is therefore an unlikely mechanism for the epidemic emergence and displacement of the predecessor lineages. Whereas some *S. pyogenes* outbreaks have been caused by strains with a hyper-encapsulation phenotype (33, 65) we are unaware of a body of epidemiologic data associating GAS epidemic outbreaks with a loss of capsule phenotype. To summarize, the global transcriptome data comparing the pre‐ and epidemic strains show that neither production of the phage encoded virulence factors SpeC and Spd1, nor lack of production of the antiphagocytic HA capsule are characteristics unique to the emergent clade 3 strains relative to the predecessor clade 1 and 2 strains and therefore do not correspond with the epidemic increase in invasive infections.

The very recent emergence of SC-3D strains is temporally associated with a single HGT event in which SC-3D strains acquired an 18-kb sequence that encodes 21 genes, including the secreted virulence proteins SpyA, a C3-like ADP-ribosyltransferase, and SpeJ, a pyrogenic exotoxin superantigen (47, 49-51). Based on near sequence identity, this 18-kb region likely was acquired from an epidemic *emm*1 clone donor. Differentially expressed genes accounted for 2% and 11% of the genome at exponential and stationary growth phases, respectively, in comparing the transcriptomes of strain MGAS26844 (clade 3) and MGAS27520 (SC-3D) (Fig. 7A, Table S3 section 3). This was the lowest number of differentially expressed genes among the four genetically representative strains studied, consistent with SC-3D strains being a recently emerged closely genetic related subset of the epidemic clade 3 strains.

### Further Transcriptome Remodeling and Epidemic Perpetuation

Discovery of significant alteration of transcriptomes caused by HGT events, and the role in emergence and dissemination of clade 3 organisms, led us to investigate the hypothesis that additional transcriptome remodeling contributed to perpetuating the *emm*89 epidemic. We tested this hypothesis by focusing on SC-3D strains, because in Finland these organisms disproportionately increased in frequency starting from 2013 (Fig. 2A, Fig. S4, Table S1). Given the relatively modest number of differentially expressed genes between MGAS26844 (clade 3) and MGAS27520 (subclade 3D), we interrogated the genome data for candidate polymorphisms that may further alter the transcriptome and potentially influence pathogen behavior. Analysis of the genome sequences of the 33 SC-3D strains found unique single amino acid replacements in gene regulators CovR (S130N) and LiaS (K214R). These polymorphisms were prevalent among the SC-3D strains, 11 strains had the CovR (S130N) change and 6 strains had the LiaS (K214R) change (Fig. S4). In contrast, none of the other 1,183 *emm*89 or 3,615 *emm*1 strains (32) studied had these polymorphisms. The branching of the strains with these mutations in the inferred phylogeny and their absence in other *S. pyogenes* strains indicates identity by descent rather than identity by independent mutation (i.e. commonality by evolutionary convergence).

Repeated recovery of clonal progeny with either the CovR (S130N) or LiaS (K214R) polymorphisms from invasive episodes has not been reported previously and thus was unexpected. Because relatively little is known about *liaS* in *S. pyogenes,* we elected to study the LiaS (K214R) polymorphism in more detail. Consistent with our altered-transcriptome hypothesis, RNAseq analysis showed that the transcriptome of strain MGAS27710 LiaS (K214R) differed from that of SC-3D LiaS wild type strain MGAS27520, including significant changes in expression of several virulence genes (data not presented). However as these two strains are not isogenic, the extent to which the altered transcription was due to the LiaS (K214R) polymorphism could not be assessed. To address this issue, we constructed a LiaS (K214R) isogenic mutant from parental strain MGAS27556 and conducted RNAseq analysis. We found that compared to the wild-type parental strain, the LiaS (K214R) isogenic mutant had 127 and 70 differentially expressed genes in exponential and stationary phase growth, respectively (Table S3 section 4). Virulence genes significantly increased in expression by the LiaS (K214R) isogenic mutant included all 9 genes of the streptolysin S biosynthesis operon *(sagABCDEFGHI)* in exponential phase and *speG* encoding streptococcal pyrogenic exotoxin G in stationary phase.

The capacity of the CovR (S130N) and LiaS (K214R) naturally occurring mutant strains to repeatedly cause serious infections means they can effectively spread between hosts, and implies that they are not attenuated in ability to survive in the upper respiratory tract, the more common *S. pyogenes* niche. Consistent with this idea, we found that the naturally occurring mutant strains had enhanced ability to survive in human saliva *ex vivo* relative to SC-3D wild type strain MGAS27520 (Fig. 8H). These results contrast with data showing that strains with other *covR/S* mutations have reduced survival in human saliva relative to wild-type strains (66).

**Fig. 8.**
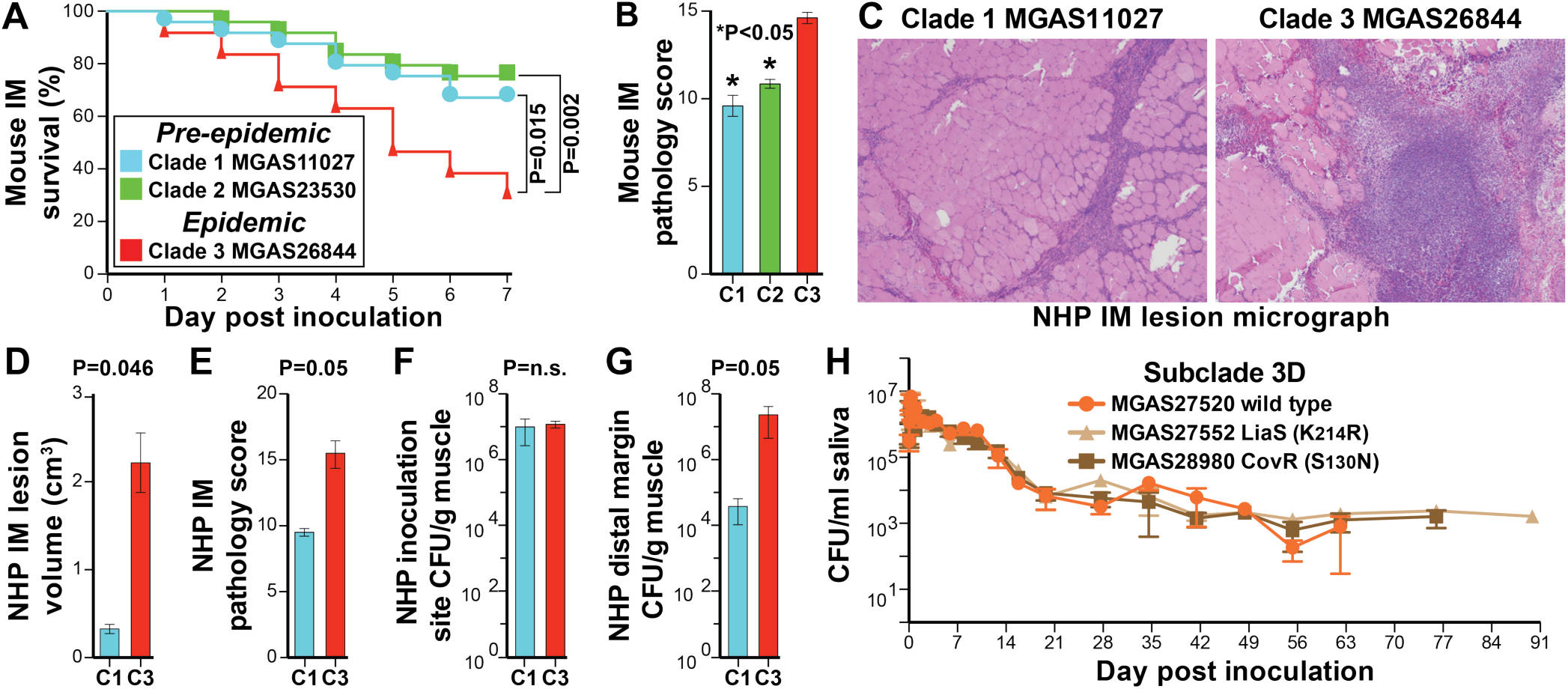
Virulence assays. (*A*) Kaplan-Meier survival curve for mice (*n*=25/strain) inoculated intramuscularly in the right hind limb with 2.5×10^8^ CFUs. The genetically representative epidemic strain (MGAS26844) was significantly more lethal than the pre-epidemic strains throughout the period of observation. The index of the strains compared in panel A applies to panels A-to-G. (*B*) Histopathology scores for muscle tissue sections as determined by pathologists blinded to the infecting strain. Illustrated is the mean (*n*=5 assessments/strain) ± SEM. P-values for panels B, D, E, F, and G were determined with the Mann-Whitney test. (*C*) Cynomolgus macaques were inoculated intramuscularly in the anterior thigh with 1.0×10^9^ CFUs/kg body mass. Shown at the same magnification are micrographs of muscle tissue sections from the site of inoculation. Epidemic strain MGAS26844 caused significantly larger lesions (panel D) with greater tissue destruction (panel E) than pre-epidemic strain MGAS11027. Although the bacterial burden was similar at the site of inoculation (panel F) it was significantly greater for the epidemic strain relative to the pre-epidemic strain at the distal margin (panel G) showing greater dissemination. (*H*) Naturally occurring variant strains MGAS28980 CovR (S130N) and MGAS27552 LiaS (K214R) viability in human saliva persisted for 2 and 4 weeks longer respectively, than that of wild type strain MGAS27520. No growth is <10 CFU/ml for a 1:10 dilution.

### Comparative Strain Virulence

The epidemiologic, comparative genomic, and transcriptome data demonstrate that clade 1, 2, and 3 organisms are genotypically and phenotypically distinct, and strongly suggest differences in virulence. To test this hypothesis, the three genetically distinct reference strains for each clade were compared in mouse and non-human primate models of necrotizing fasciitis (NF) (67-69). Epidemic clade 3 reference strain MGAS26844 was significantly more lethal and caused significantly greater tissue damage in the mouse NF infection model than the two pre-epidemic reference strains (Fig. 8A-B). Moreover, relative to clade 1 reference strain MGAS11027, epidemic clade 3 strain MGAS26844 caused significantly larger lesions with greater tissue damage in a nonhuman primate model of NF (Fig. 8C-E).

## Concluding Comment

We have used *S. pyogenes* as a model pathogen for studying the evolutionary genomics of epidemic disease and the molecular basis of bacterial pathogenesis. The organism is a strict human pathogen, causes abundant infections worldwide and has a relatively small genome (~1.8 Mb). In addition to its propensity to cause epidemic waves, the availability of comprehensive, population-based strain collections from many countries, coupled with the fact that humans are its only natural host, means that the history of underlying events that generate genomic diversity is not obscured by molecular processes occurring in non-human hosts or environmental reservoir. These factors afford considerable advantages to *S. pyogenes* as a model system compared to many other pathogenic bacteria such as *E. coli, S. enterica,* and *S. aureus.*

The primary goal of our study was to determine if genomic changes linked with the origin and perpetuation of human epidemic disease have remodeled global gene expression and altered virulence in the model pathogen *S. pyogenes.* We were especially interested in determining the effect, if any, of horizontally acquired genome segments on global gene expression and virulence of the progeny strains. Despite the importance of bacterial pathogens in human and veterinary health, remarkably few studies have addressed how transcriptome remodeling contributes to the origin and perpetuation of epidemics. Zhou et al. (26) studied diversity in 149 genomes of *S. enterica* serovar Paratyphi A and used the resulting data to speculate that most recent increases in frequencies of bacterial diseases are due to environmental changes rather than the novel evolution of pathogenic bacteria. In essence, it was suggested that many epidemics and pandemics of bacterial disease in humans did not involve recent evolution of particularly virulent organisms, but instead reflected chance environmental events. A similar conclusion was reached in studies of other pathogens, for example *Yersinia pestis*, *S. enterica* serovar Agona, *Mycobacterium tuberculosis*, *Mycobacterium leprae*, and *Shigella sonnei* (17, 25).

Although this may be the case for some pathogens, based on the full-genome data from 4,815 strains, human patient information (33), analysis of isogenic mutant strains, RNAseq studies, and experimental animal infection, we arrive at a fundamentally different conclusion for *emm*89 and *emm*1 *S. pyogenes*, organisms that have caused epidemics involving tens of millions of human infections in the last 30 years. In particular, our results unambiguously show that newly emerged clones causing epidemic disease are more virulent than previously circulating precursor organisms. For clarity, we consider all steps in pathogen-host interaction to potentially contribute to the virulence phenotype, including survival and proliferation after initial contact with the host through invasion of deeper tissues and spread to new hosts. Conclusions about molecular pathogenesis and virulence based solely or predominantly on population genomic analyses of a convenience sample of strains and resulting inferences are not likely to fully reflect the biology of pathogen and host interaction. This issue may be especially problematic if only one or a few nucleotide changes significantly alter virulence.

We believe that our findings have important implications for bacterial pathogens that must successfully circumvent host defenses, both at the individual and population level. Our analysis demonstrated that among the various *emm*89 clades and subclades, considerable variation exists among global transcriptomes, both in the spectrum of genes expressed and their magnitude of expression. This means that in essence many different antigen, toxin and virulence factor profiles can and are being displayed to host populations as a function of individual strain genotype, and not necessarily by *emm-* type. In the absence of one or a small number of conserved antigens mediating protective immunity, regardless of the microbe, significant variation in antigen repertoire has implications for vaccine research, formulation, and deployment.

Many elegant studies of the population genomics of bacterial pathogens have been published over the last decade (4, 5, 10-18, 23, 25, 26, 32, 70-74). There is a small but emerging literature bearing on the impact of regulatory plasticity in bacterial evolution and fitness (75-79). However, there has been very little work designed to integrate microbial population genomics, molecular pathogenesis processes, microbial emergence, transcriptome remodeling, and virulence. Our findings suggest this could be a fruitful area of research for other microbial pathogens. The resulting data are likely to have significant implications for understanding bacterial epidemics and translational research efforts to blunt their detrimental effects.

## Materials and Methods

Further details of materials and methods are described in *SI Materials and Methods.*

### Bacterial Strains

We studied 1,200 GAS *emm*89 strains, including 1,198 strains causing invasive infections and two from pharyngitis patients (Table S1). The vast majority of the strains (*n*=1,178) were collected as part of comprehensive population-based public health surveillance of GAS invasive infections conducted in the United States, Finland, and Iceland between 1995 and 2014. The remaining *emm*89 strains were recovered from invasive disease cases in Ontario, Canada and a pharyngitis case in Italy. A subset of this population has been previously studied and preliminary genetic findings presented (35, 48).

### Genome Sequencing

Isolation of chromosomal DNA, generation of paired-end libraries, and multiplexed sequencing were accomplished as described previously (32, 35) using Illumina (San Diego, CA) instruments (HiSeq2500, MiSeq, NextSeq). Whole genome sequencing data for the 1200 isolates studied were deposited in the NCBI Sequence Read Archive under accession number SRP059971.

### Reference Genome Assembly, Annotation, and Polymorphism Discovery

The bioinformatics tools used for assembling and annotating the reference genomes, and for identifying and analyzing polymorphisms in the population studied are described in *SI Materials and Methods.* Complete genome sequences for strains MGAS11027, MGAS23530, and MGAS27061 were deposited in NCBI GenBank database under accession numbers CP013838, CP013839, and CP013840 respectively. MGAS11027, MGAS23530, and MGAS27061 were deposited in the BEIR strain repository under accession numbers NR-33707, NR-33706, and NR-50285 respectively.

### Phylogenetic Inference and Population Structure

The bioinformatics tools used for sequence alignments, detection and filtering of HGT polymorphisms, clustering, phylogenetic inference and analysis of the population structure are described in *SI Materials and Methods.*

### Gene Content and Mobile Genetic Element Analysis

The known GAS pangenome core and accessory gene content was determined based on 30 complete genomes of 18 different emm-types (Table S2) as described in *SI Materials and Methods.* Among the 53,336 CDSs of the 30 genomes, PanOCT identified 3,338 ortholog clusters which was culled by BLAST reciprocal-best-hit to 2,835 on the basis of no two clusters sharing >95% amino acid identity. A GAS pseudo-pangenome sequence of ~3 Mbp was generated by concatenating onto the *emm*89 MGAS23530 reference genome all accessory gene content not already present in the genome, starting with *emm*89 strains MGAS11027 and MGAS27061, and then the remaining 27 genomes by emm-type (i.e. *emm*1, 2, 3, etc.). Based on mapping of the *emm*89 reference genome sequencing reads to the GAS-30 pangenome, a RPKM (reads per kilobase of transcript per million reads mapped) value of >50 corresponded with gene presence. A phage was called present if a minimum of 80 percent of its gene content represented in the GAS-30 pangenome was determined present. Reads not mapping to the GAS-30 pangenome were assembled *de novo* using SPAdes. Resultant contigs greater than 100 nucleotides were queried against the NCBI nonredundant database using BLAST to determine their nature.

### Construction of Isogenic Mutant Strains

The construction of the *liaS* isogenic mutant strain was accomplished by allelic exchange as previously described (35). Briefly, MGAS27556 LiaS (K214R) was generated by introducing the *liaS* A641G SNP into wild-type strain MGAS27556, using DNA amplified from strain MGAS27552, a clinical isolate with a naturally occurring *liaS* A641G SNP (i.e. LiaS K214R substitution) as template. Successful introduction of the desired SNP and the absence of spontaneous spurious mutations, were confirmed in candidate isogenic mutants by whole genome sequencing. Primers, plasmids, and restriction enzymes used in the construction are listed in *SI Materials and Methods.*

### Transcriptome Sequencing and Expression Analysis

Whole genome transcriptional analysis was conducted for representative strains using RNAseq as previously described with minor modifications (35, 80). Briefly, RNA was isolated from triplicate cultures grown in THY. Multiplexed libraries were single-end sequenced (50 bp) to high depth (~10 million reads/sample) with an Illumina HiSeq2500 instrument. RNAseq reads were mapped to the genome of the most closely related *emm*89 reference strain (for example, clade 3 strains were mapped to the genome of reference strain MGAS27061). Use of multiple reference sequences was critical, as the use of a single common reference did not permit accurate quantitative read mapping to the divergent sequences in the regions of HGT. RNAseq data was normalized and genes statistically differently expressed following Benjamini-Hochberg correction at a minimum 1.5 fold change in mean transcript level were identified using bioinformatics tools provided in *SI Materials and Methods.* RNAseq transcriptome data were deposited in the NCBI Gene Expression Omnibus database under accession number GSE76816.

### Experimental Animal Infections

The virulence of serotype *emm*89 reference strains MGAS11027, MGAS23530 and MGAS26844 was assessed using mouse and non-human primate models of necrotizing fasciitis (32, 67-69). These strains have a wild-type (i.e. the most commonly occurring) allele for all major transcription regulators, including *covR/S, ropB,* and *mga.* All animal experiments were approved by the Institutional Animal Care and Use Committee of Houston Methodist Research Institute.

## ACKNOWLEDGMENTS

We thank FiRe - the Finnish Study Group for Antimicrobial Resistance, Chris A. Van Beneden, Bernard Beall and the Active Bacterial Core surveillance (ABCs) of the Center for Disease Control and Prevention’s (CDC) Emerging Infections Programs network; Kathryn Stockbauer and Helen Chifotides for critical comment and editorial assistance; and Hanne-Leena Hyyrylainen, Kai Puhakainen, and Francesca Latronico for microbiological and epidemiological assistance. This project was supported in part by the Fondren Foundation, Houston Methodist Hospital; the Academy of Finland (grant 255636); and by the European Society of Clinical Microbiology and Infectious Diseases Training Fellowship 2011 and the Federation of European Societies of Microbiology Research Fellowship 2011-1 awarded to M.C.D.L.

## Supplemental File Legends

**Text S1.** Supplemental Materials and Methods

**Fig. S1.** Atlases for the three *emm*89 reference genomes. Shown from the outer-most (first) ring to the inner-most (twelveth) ring are the following. 1) Megabase-pairs (black); 2) Gene or operon landmarks; 3 and 4) coding sequences on the forward strand (light-blue) and reverse strand (dark-blue); 5, 7, and 9) BLAST nucleotide sequence comparison with the genomes indicated in the respective indexes; 6, 8, and 10) distribution of SNPs for the genomes indicated in the respective indexes; 11) G+C relative to the mean; and 12) GC skew. BLAST nucleotide sequence comparisons were made between the genomes of the clade 1, 2, and 3, reference strains and with a de novo assembly of strain MGAS27450 the most phylogenetically distant *emm*89 outlier strain.

**Fig. S2.** Genetic relationships among *emm*89 reference strain, with *emm*1 reference strain SF370 used as the rooting outgroup. Shown are genetic relationships among the three *emm*89 reference strains and the seven outlier strains using *emm*1 reference strain SF370 as an outgroup. Relationships were inferred based on 26,371 core SNPs by neighbor-network splits-decomposition. The sequence of branching of the three numerically dominant *emm*89 primary clades along the evolutionary path leading to the contemporary epidemic *emm*89 strains is clade1 (MGAS11027), followed by clade 2 (MGAS23530) and then epidemic clade 3 (MGAS27061).

**Fig. S3.** Potential horizontal gene transfer (HGT) region donors. Shown for each of the predicted recombination blocks (RB) separating the clades, are the genetic relationships among the three *emm*89 clade reference strains and 39 strains of 18 other *emm* types for which there are complete genome sequences publically available as of July 10, 2015. Sequences flanking the predicted recombination blocks in the strain MGAS23530 genome were used to define the corresponding regions in the other strains using blastn. The sequences corresponding to the predicted recombination blocks among all 42 strains were aligned using MAFFT, and relationships were inferred by neighbor-network split-decomposition using SplitsTree. The length of the recombination block and locus tags of the genes involved, are listed relative to strain MGAS23530. Of note, seven of the eight recombination blocks separating all 359 clade 1 strains from all 78 clade 2 strains share a more recent common ancestor with *emm*2 reference strain MGAS10270 than with reference strains of any of the other emm types used in this comparison.

**Fig. S4.** Genetic relationships among *emm*89 subclade 3D strains. Shown are genetic relationships among the 33 subclade 3D strains using clade 3 reference strain MGAS27061 as an outgroup. Relationships were inferred based on 157 core SNPs by neighbor-joining using SplitsTree. All subclade 3D strains differ from all progenitor clade 3 strains by an 18 kb region of HGT involving the virulence factors SpyA and SpeJ (Table 1, recombination block 15). To constrain the inference primarily to vertically inherited SNPs, SNPs within putative regions of HGT were identified and filtered out using GUBBINs. The 11 strains with the CovR (S130N) substitution branch together, indicating inheritance by descent. Similarly, all but one of the 6 strains with the LiaS (K214R) substitution branch together again indicating inheritance by descent. We attribute the single LiaS (K214R) strain not branching with the others as likely being due to a few scant horizontally acquired polymorphisms that were insufficient to statistically significantly elevate the SNP density and therefore were not detected/excluded by GUBBINs.

**Fig. S5.** Comparison of phages 11027.1 and 27061.1. Shown above is a percent identity plot and below is a dot matrix alignment. The phages are similar over the 5’ first ~13 kb encoding the integrase, replication and lytic/lysogenic regulatory genes, diverge over most of the central portions encoding head and tail coat proteins, and then are similar again over the 3’ last ~3 kb encoding the secreted virulence factors streptococcal pyrogenic exotoxin C (SpeC) superantigen and the streptococcal phage DNase 1 (Spd1). The divergence in sequence between 11027.1 and 27061.1, means that 27061.1 did not evolve from 11027.1 through a simple single deletion event. Despite being integrated at the same genomic locus and encoding the same virulence factors, they are distinct mosaic phages.

**Fig. S6.** *hasABC* promoter variants. (*A*) Illustrated are *hasABC* promoter pattern variants identified among the *emm*89 clade 1 and and 2 strains. Patterns A and B account for 99% of the strains. Pattern B has a 38 bp deletion relative to pattern A, which eliminates a putative Rho independent terminator. In M1 strain MGAS2221, deletion of this terminator results in release of *hasABC* from CovR repression resulting in enhanced capsule production. (*B*) Distribution of *hasABC* promoter variants among the clade 1 and 2 strains. (*C*) Distribution of *hasABC* promoter variants among the clade 1 and 2 strains. Illustrated is genetic relationships among the *emm*89 clade 1 and 2 strains inferred by neighbor-joining based on 5,663 core SNPs filtered using GUBBINs to exclude regions of horizontal gene transfer. Strains are colored by promoter variant as indicated in the index. Clade 1 strains are a nearly equal mix of pattern A (weak/repressed) and pattern B (strong/derepressed) promoter variants, whereas the vast majority of clade 2 strains are pattern A. These findings are consistent with the significantly greater level of *hasABC* transcripts for clade 1 strain MGAS11027 relative to clade 2 strain MGAS23530 determined by RNAseq.

**Table S1.** Strains and Characteristics

**Table S2.** *Streptococcus pyogenes* Complete Genome Sequences

**Table S3.** RNAseq Transcriptome Analyses

